# Outbreak of invasive wound mucormycosis in a burn unit due to multiple strains of *Mucor circinelloides* f. *circinelloides* resolved by whole genome sequencing

**DOI:** 10.1101/233049

**Authors:** Dea Garcia-Hermoso, Alexis Criscuolo, Soo chan Lee, Matthieu Legrand, Marc Chaouat, Blandine Denis, Matthieu Lafaurie, Martine Rouveau, Charles Soler, Jean-Vivien Schaal, Maurice Mimoun, Alexandre Mebazaa, Joseph Heitman, Françoise Dromer, Sylvain Brisse, Stéphane Bretagne, Alexandre Alanio

## Abstract

*Mucorales* are ubiquitous environmental molds responsible for mucormycosis in diabetic, immunocompromised, and severely burned patients. Small outbreaks of invasive wound mucormycosis (IWM) have already been reported in burn units without extensive microbiological investigations. We faced an outbreak of IWM in our center and investigated the clinical isolates with whole genome sequencing (WGS) analysis.

We analyzed *M. circinelloides* isolates from patients in our burn unit (BU1) together with non-outbreak isolates from burn unit 2 (BU2, Paris area) and from France over a two-year period (2013-2015). For each isolate, WGS and a *de novo* genome assembly was performed from read data extracted from the aligned contig sequences of the reference genome (1006PhL).

A total of 21 isolates were sequenced including 14 isolates from six BU1 patients. Phylogenetic classification showed that the clinical isolates clustered in four highly divergent clades. Clade1 contained at least one of the strains from the six epidemiologically-linked BU1 patients. The clinical isolates seemed specific to each patient. Two patients were infected with more than two strains from different clades suggesting that an environmental reservoir of clonally unrelated isolates was the source of contamination. Only two patients shared one strain in BU1, suggesting direct transmission or contamination with the same environmental source.

WGS coupled with precise epidemiological data and analysis of several isolates per patients revealed in our study a complex situation with both potential cross-transmission and multiple contaminations with a heterogeneous pool of strains from a cryptic environmental reservoir.

**Importance:** Invasive wound mucormycosis (IWM) is a severe infection due to the environmental molds belonging to the order Mucorales. Severely burned patients are particularly at risk for IWM. Here, we used Whole Genome Sequencing (WGS) analysis to resolve an outbreak of IWM due to *Mucor circinelloides* that occurred in our hospital (BU1). We sequenced 21 clinical isolates, including 14 from BU1 and 7 unrelated isolates, and compared them to the reference genome (1006PhL). This analysis revealed that the outbreak was mainly due to multiple strains that seemed patient-specific, suggesting that the patients were more likely infected from a pool of diverse strains from the environment rather than from direct transmission between the patients. This study revealed the complexity of a *Mucorales* outbreak in the settings of IWM in burn patients, which has been highlighted based on whole genome sequencing and careful sampling.

## Introduction

Mucormycosis is a rare and life-threatening infection caused by *Mucorales* belonging to the subphylum *Mucoromycotina* (1). These molds are ubiquitously distributed in the environment and mostly disseminated through airborne spores, which can be considered as infective propagules responsible mainly for respiratory (lung and sinuses), wound, and skin infections (2).

Patients at risk for mucormycosis are immunocompromised (hematological malignancies, hematopoietic stem cell transplantation, solid organ transplant, steroid therapy), or have diabetes mellitus, deferoxamine treatment, trauma or severe burns (2–4). Among skin-related infections, contaminated materials (Elastoplast bandages, tape, tongue depressors, ostomy bags, linens) have been implicated as the cause of local or disseminated infections in patients with various underlying diseases (5). Specifically, in burn patients, invasive wound mucormycosis (IWM) has been reported in both small series (6) and epidemiological surveys (7–10).

Over the past 10 years, outbreaks of mucormycosis have been increasingly reported in various environments. Indeed, outbreaks in animals including chickens, sheep and frogs have been described (11–13). In humans, outbreak cases in the hospital before 2008 were reviewed by Antoniadou in 2009 (14). The author found 12 reported outbreaks and two pseudoepidemics of cases since 1977. Outbreaks of mucormycosis have been reported in the USA, UK and Europe (14). Since 2008, outbreaks or clustered cases have been reported after the tornadoes in Joplin, Missouri (13 patients) (15, 16), in the ICU in France (three patients) (17), in adults (six patients) (18) or infants (five patients) (19) exposed to contaminated linens in the US, in infants in Egypt (five patients) (20), and in patients undergoing arthroscopy in Argentina (40 patients) (21). More specifically, in a Belgian burn unit, Christiaens *et al.* described an outbreak of *Lichtheimia corymbifera* associated with non-sterile elastoplast bandage contamination in seven burn patients including five with infection and two with colonization (22). In this study, the authors did not have evidence for the genotypic relatedness of the strains between patients and material strains. In addition, a large outbreak due to yogurt contamination in the US responsible for digestive symptoms (nausea, cramps, vomiting, and diarrhea) in about 300 individuals has been described recently (23). In this study, phylogenetic analysis and whole genome sequence (WGS) analysis yielded new information on the genetic structure of *Mucor circinelloides* species complex, which may include 3 related but distinct species currently recognized as *Mucor circinelloides f. circinelloides, lusitanicus,* and *griseocyanus* (23, 24).

Between 2013 to 2015, we faced an outbreak of proven IWM due to *M. circinelloides* f. *circinelloides* in a burn unit (BU) in the Saint-Louis Hospital (SLS), Paris, France involving six patients raising the hypothesis of a common source of contamination. Over the same period of time, 4 additional cases that occurred in a burn unit (BU2) of another hospital (PER, Clamart, France) in the Paris suburb were notified the National Reference Center for Invasive Mycoses and Antifungals (NRCMA).

Our aim was to clarify the origin/source of infection in both BU1 and BU2. In the absence of genotyping markers for this organism, WGS analysis was performed on 21 isolates (14 from the outbreaks and 7 unrelated) to investigate the links between clinical isolates, understand the epidemiology of the outbreak, and identify and eliminate the source of the infections.

## RESULTS

### Clinical and microbiological investigation

Three patients (P03, P04, P05) developed proven IWM due to *M. circinelloides* f. *circinelloides* within 18 days in BU1 (between August 18th and September 5^th^, 2014), and subsequently died from these infections (Table 1). The outbreak was suspected when a positive culture was observed in P04 (11 days after the first positive sample in P03). Sequential samples from the wounds were prospectively obtained starting with P03. A few months later (114 days), another patient (P06) developed proven IWM and *M. circinelloides* f. *circinelloides* was also involved (Fig. 1). We retrospectively noticed that isolates from the same species have already been identified in 2013 from two patients on BU1 (P01 with no infection, and P02 with proven IWM). A total of six patients were exposed (P01, P07) and/or infected with this species (P02 to P06) in BU1, with P01 as the putative index case. Infection control measures were implemented locally to avoid potential nosocomial transmission to other patients of the unit. More than 30 environmental samples were cultured. All were negative. DNA amplified with the Mucor/Rhizopus PCR test (25) was detected only in the Bair Hugger filters that were used during the hospitalization of P03, P04 and P05.

**Table 1:**
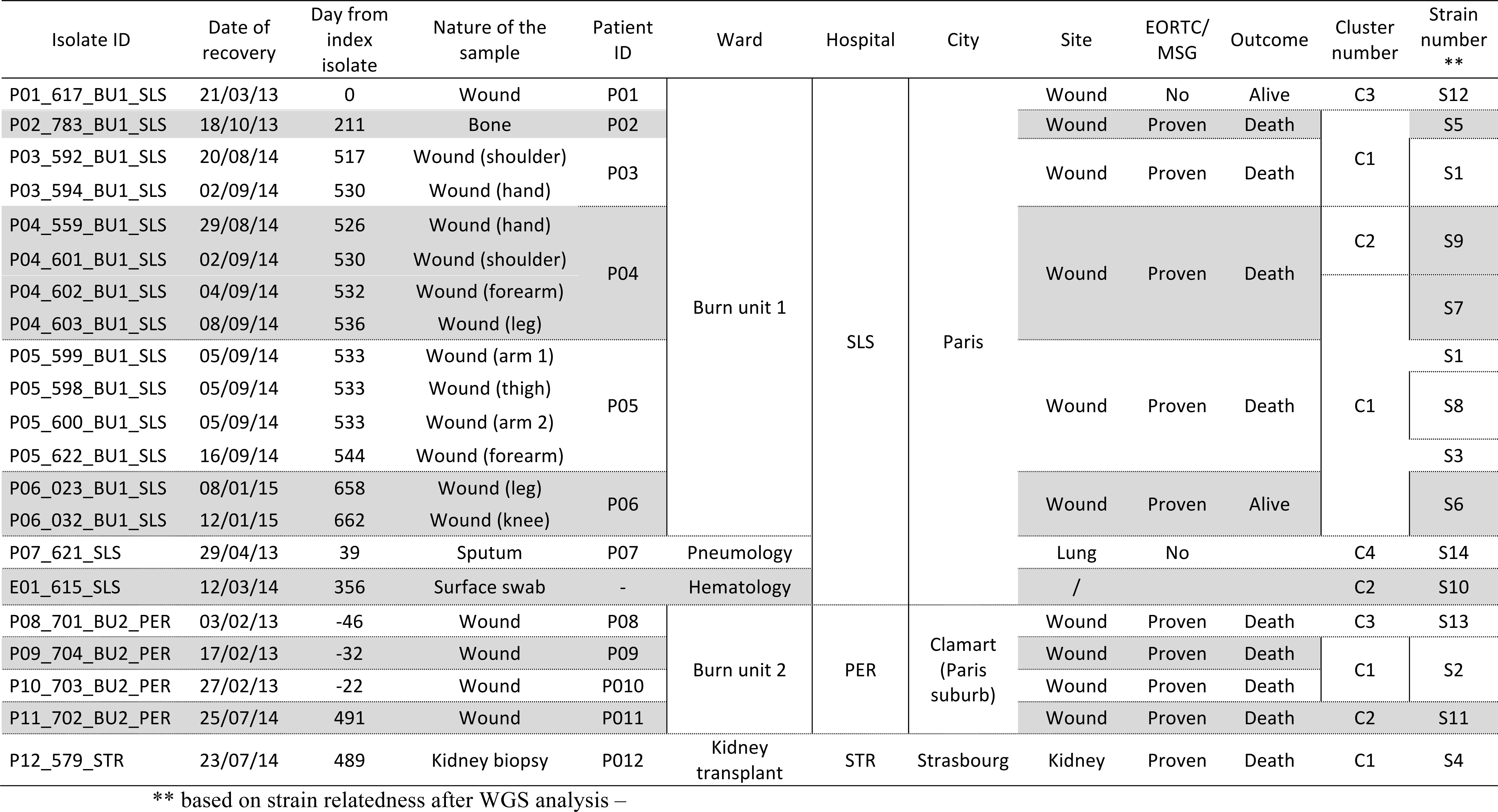
Isolates sequenced in this study

**Fig. 1.**
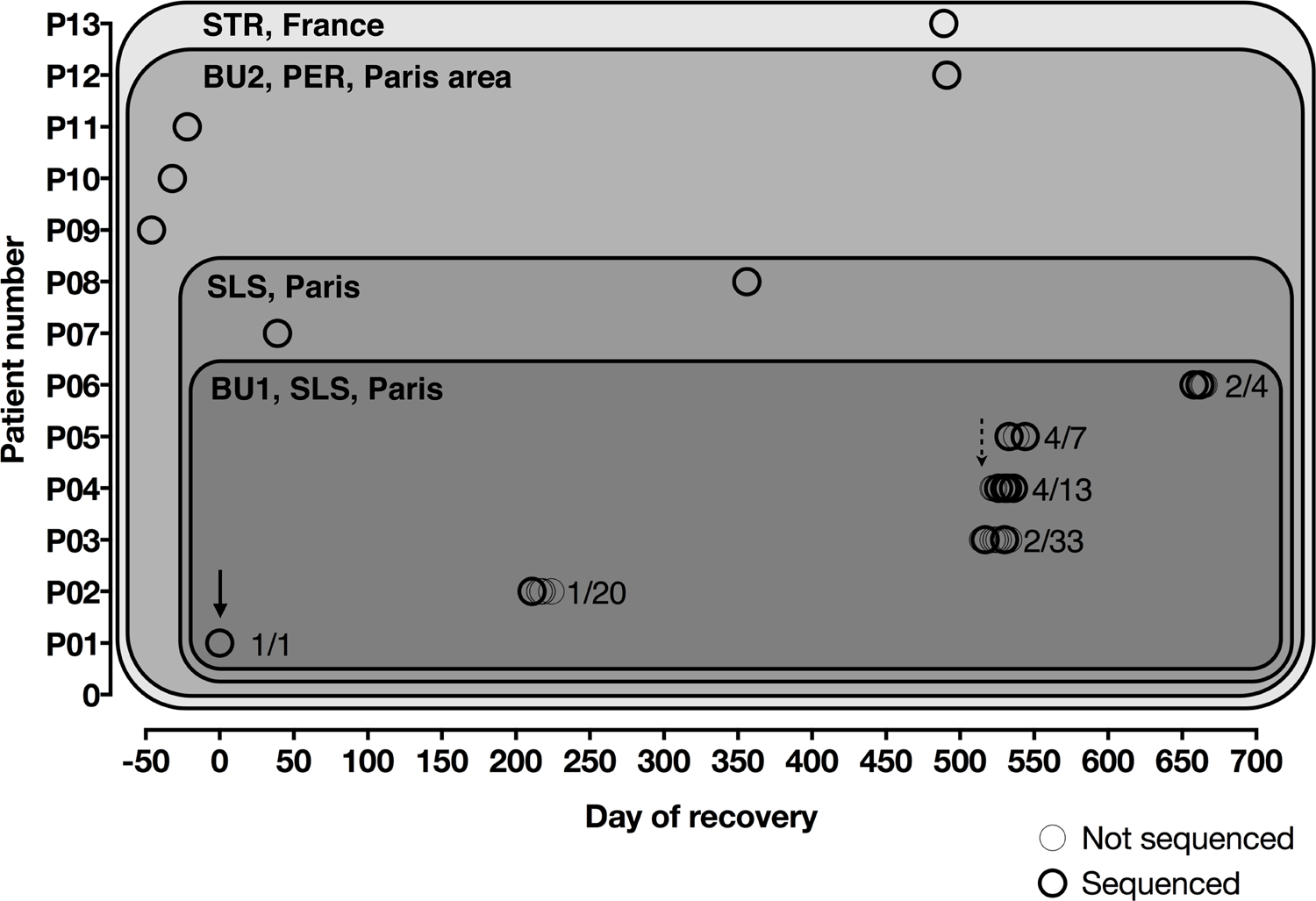
Epidemiological map of thirteen patients whose isolates were selected in our study. Isolates from several geographic areas in France (from dark to light gray) have been studied: burn unit 1 of Hospital 1, Paris France; wards of SLS, Paris, France; Burn unit 2 of PER in Paris area, France and eastern France (one isolate from STR). Isolates were prospectively collected for patients P03 to P06. Analyzed isolates are depicted in dark open circles. Index case of burn unit 1 (full line arrow) was thought to be P01 and outbreak have been recognized after P04 got infected (dashed arrow)

To prevent transmission to other patients, we needed to investigate whether these strains were clonal, and needed unrelated isolates from other geographical areas. The additional cases corresponded to an outbreak in BU2 (PER) involving four patients with IWM, as well as one case of proven invasive mucormycosis with kidney invasion in a transplant recipient in STR that was notified to the NRCMA.

Overall, the 12 patients (21 clinical isolates) included 10 cases identified in two burn units (Table 1, Fig. 1).

### Phylogenetic analyses of three loci

ITS, D1/D2, and RPB1 sequences were analyzed as separate (data not shown) and combined datasets. The topology of the multi-locus dataset with 3 methods (NJ, ML and Bayesian inference) was comparable between the individual trees of the three genes analyzed. Four clades (Fig. 2), respectively denoted C1 (14 isolates including 11 from BU1), C2 (four including two from BU1), C3 (two strains in addition to the reference strain including one from BU1) and C4 (one isolate) were identified from the analysis of the combined dataset that yielded a significant support (≥95% bootstrap for NJ and ML; 1.0 for Bayesian inference). Isolates recovered from BU1 were distributed in three clades (C2, C3, C4). All patients from BU1 had at least one isolate included in C1.

**Fig. 2.**
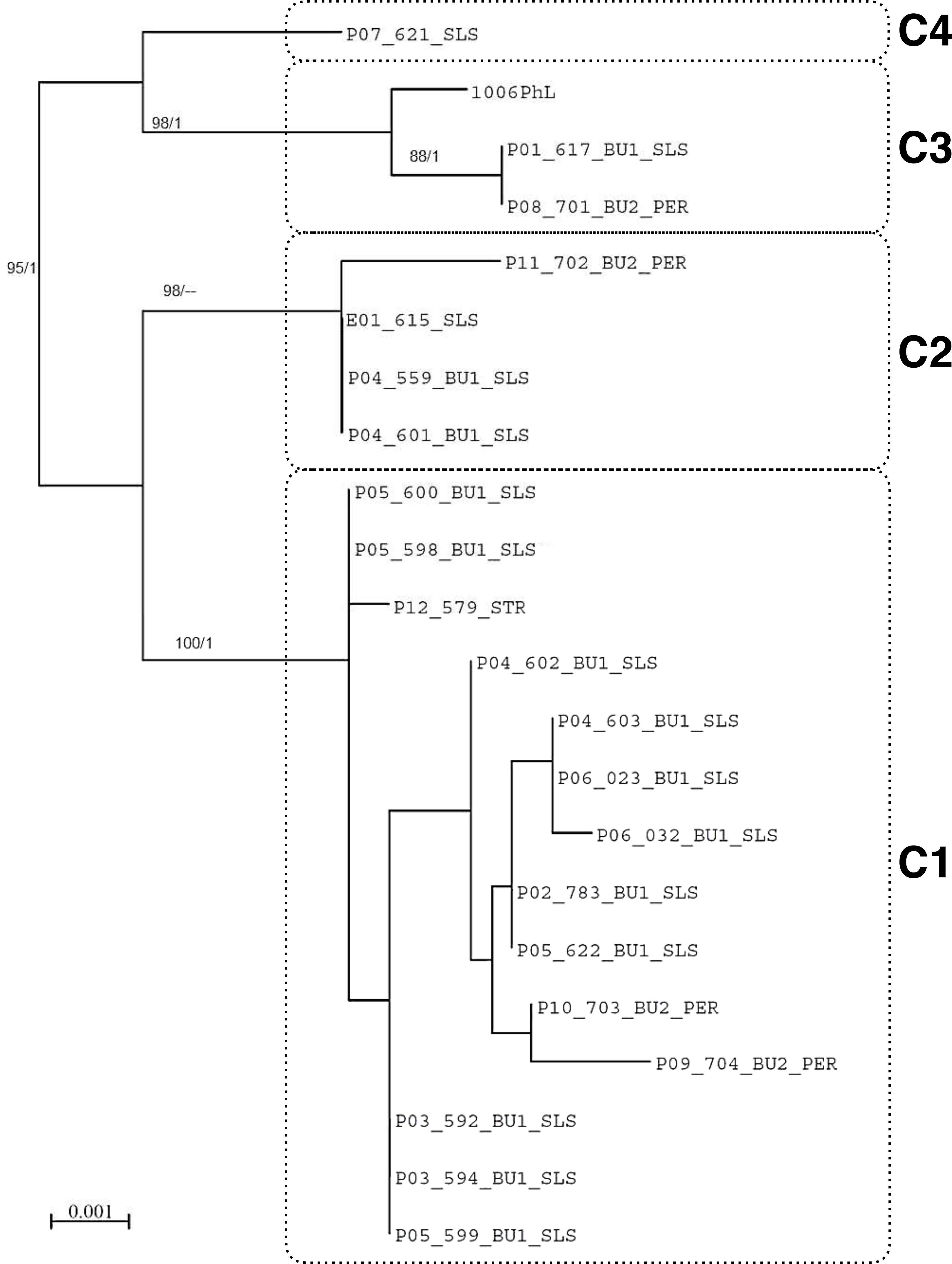
PhyML tree constructed Maximum Likelihood tree inferred from concatenated 3 loci dataset (ITS, 28S, and RPB1). Bootstrap support values from PhyML greater than 70% (left) and Bayesian posterior probability >0.80 (right) are shown at the nodes. The 21 clinical isolates are grouped in clades C1 to C4. The scale bar indicates 0.001 nucleotide substitutions per character.

### Whole genome analysis

To better resolve the diversity of the strains within the four clades, and because no further genotyping methods existed for this organism, whole genome sequencing was performed. Because the biology and the genetics of this organism is poorly understood, we first checked the reproducibility of the sequencing process and the stability of the genome, to be able to define genetically-identical strains.

#### Establishing the genetic threshold to determine genetically-identical strains

For the three strains isolated from single spore colonies (i.e. P05_600_BU1_SLS, P04_603_BU1_SLS and P03_594_BU1_SLS), the pairwise evolutionary distance was estimated between *de novo* assemblies of the parental and the single spore colony (i.e. 0.00035, 0.00044 and 0.00029, respectively). This information gave us the expected distances between pairs of genomes arising from identical strains and independently sequenced distinct isolates. The largest of the three distances (i.e. 0.00044) was therefore selected as a cutoff below which two compared isolate genomes were defined as arising from the same strain.

#### Experimental investigation of the potential genetic drift of M. circinelloides f. circinelloides

Experimental investigation of the 1006PhL genome upon iterative sub-culturing on agar (n=3) and three passages in mice (n=3) revealed no acquisition of SNPs during this process, suggesting that the genome of *M. circinelloides* f. *circinelloides* was stable upon iterative passages.

#### Whole genome phylogenetic classification of the 21 clinical isolates

A phylogenetic classification of the whole genome of the 21 isolates was then performed (Fig. 3). This phylogenetic tree allows classification of the genomes in four main clades corresponding exactly to the same clades (C1 to C4) described based on the analysis of three loci (Fig. 2). Clades C2, C3, and C4 contained isolates that are clearly distinct from those inside clade C1, e.g. estimated pairwise evolutionary distances between isolates from C1 and those inside C2, C3, and C4 are 0.0187, 0.0385, and 0.0390 on average, respectively, whereas C1 pairwise intra-distance is 0.0014 on average.

**Fig. 3.**
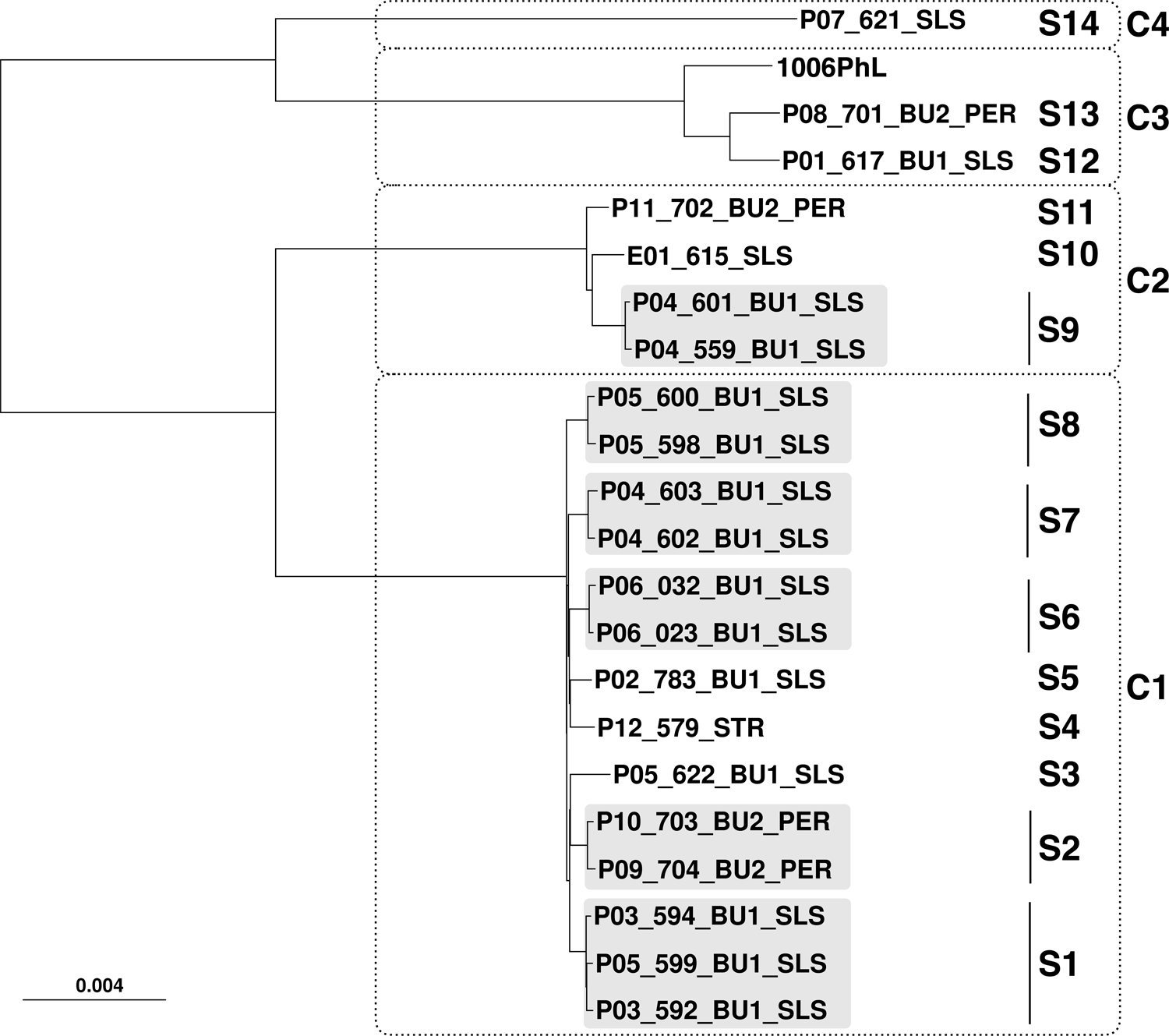
Minimum Evolution phylogenetic tree of the whole genome of 21 clinical isolates and the 1006PhL reference strain. The 21 isolates are grouped in different strains clustered in clades C1 to C4. Clinical isolates belonging to the same strain are highlighted in light grey. The scale bar indicates 0.04 nucleotide substitutions per character.

Furthermore, increased resolution of WGS allowed to robustly identify strains and understand which clinical isolates belong to which strain. Indeed, the 21 investigated isolates and the reference 1006PhL could be partitioned into 14 distinct strains (S1 to S14; Fig. 3, Table 1).

### Outbreak dynamics

In BU1, the isolate of the potential index case P01 (P01_617_BU1_SLS, S12) was different from the isolates subsequently recovered in BU1. The isolate from P02, corresponding to the specific strain S5 clustered in C1 together with isolates from P03, P04, P05, and P06. P03 and P06 were also infected with two isolates from S1 (#594 and #592) and S6 (#032 and #023), respectively. P04 was infected over 10 days with two strains S9 and S7 belonging to C2 (isolates #601 and #559) and C1 (isolates #602 and #603), respectively. P05 was infected over 11 days with isolates belonging to three strains from C1, S8 (isolates #598 and #600) and S3 (#622) and S1 (#599). Indeed, P04 and P05 had mixed infections during the course of their disease, suggesting initial contamination with a mixture of strains, this latter strain also recovered in P03. This suggests cross contamination or common infection in P3 and P5. This have also been observed in BU2, where two patients (P09 and P10) shared the strain S2 (#703 and #704) that clustered in C1. Two patients (P11 and P08) were also infected with strains that belong to C2 and C3, respectively. The environmental (#615) and the colonization (#621) isolates from our hospital clustered in C2 and C4 respectively. The patient from Eastern France clustered in C1 (S4) but with a specific and different strain than the other C1 strains, as expected for a geographically unrelated infection.

## DISCUSSION

Because isolation of Mucorales is rare in the hospital, the observation of the same species in two independent samples/patients has long been considered as a sufficient criterion to suspect and assess transmission or common contamination. Here, we investigated further outbreak-related and-unrelated isolates based on WGS analysis. To our knowledge, this is the first time WGS analysis has been utilized to resolve an outbreak of invasive *Mucorales* infection in the context of nosocomial acquisition. Because WGS was applied for the first time in this setting, we first sought to evaluate reproducibility of the sequencing process and intra-culture variation/stability during in vivo passage. As the genome of three selected strains was sequenced and assembled twice, we were able to compare contig sets belonging *a priori* to the same strain. This method led to the definition of a pairwise distance cutoff. Therefore, if two genome sequences belonging to different isolates have a pairwise distance below this cutoff, it was inferred *a posteriori* that the two isolates corresponded to the same strain. This cut-off should vary as a function of the organism, the method of sequencing, the bioinformatics pipeline and the pathophysiology of the disease, reinforcing that such data should be obtained each time an investigation of an outbreak due to rare organisms is undertaken.

Of note, using pairwise evolutionary distances could be considered as a fast but accurate alternative to the well-known Average Nucleotide Identity (ANI) approach to compare genomes (26, 27) because both were shown to be linearly correlated (28).

Contrary to the initial hypothesis of a single strain transmission in BU1, we observed that all of the patients from the BU1 outbreak (P01 to P06) were infected by different strains. Surprisingly, our data revealed that each strain was patient-specific in BU1, except for S1, suggesting that the outbreak in BU1 was due to multiple strains present in and acquired from a local environmental “reservoir” containing clonally unrelated isolates. This hypothesis is reinforced by two patients (P04 and P05) with IWM coinfected by more than 2 genetically distinct strains. Another hypothesis is that the patients could have been exposed to specific strains or a mixture of strains before arriving in BU1. However, a delay between admission and the first positive culture was 16 days (median), making the hypothesis that exposure occurred in the environment of BU1 more likely. In the settings of severe burns where invasive fungi do sporulate on the wounds, transmission by air or by the hands of healthcare workers to other patients are both possible. A major point to emphasize is that only two patients from BU1 (P03 and P05) shared the same strain S1. This feature has also been observed in BU2 suggesting that transmission between patients is possible. However, this does not rule out the hypothesis of a contamination by the same strain from the environment.

Environmental investigation of the outbreak in BU1 failed to identify the source of infection using culture of multiple samples from the environment, as well as by PCR. DNA amplified with the Mucor/Rhizopus PCR (25) was detected only in the Bair Hugger filters that were used during the hospitalization of the IWM patients P03, P04 and P05. Despite the negative result of this investigation, it is likely that the contamination came from a local source because this has already been described with linens or 12lastoplast in burn units (5, 18, 19).

Our findings about the genetic structure of *M. circinelloides* f. *circinelloides* are reminiscent of the WGS investigations of the *Apophysomyces spp.* outbreak in Joplin (16), or of the *S. clavata* outbreak in France (29), which revealed that several genetic groups can be responsible for infections over the same period of time. In our case, in a given restricted area (BU1), we identified a large diversity of isolates responsible for IWM and were not able to find isolates belonging to unique strains recovered in different places.

At the other end of this spectrum, for *Apophysomyces trapeziformis* several genetically-identical isolates were recovered in different places at a distance of several miles (16). In the case of *E. rostratum,* all outbreak isolates have closely-related genomes suggesting that a unique strain was responsible for the outbreak (30). These differences could be explained by a completely different pathophysiology of the disease and mode of fungal transmission between these fungal organisms.

This outbreak is illustrative of the importance of the sampling strategy. Repeated sampling is paramount, as is avoiding the assumption that one patient should harbor only one strain. Indeed, in our case two patients were infected by mixtures of strains concomitantly, as already described for cryptococcosis (31). Mixed infections and the impact of genetic heterogeneity of isolates from single patients on the interpretation of transmission chains is an emerging theme in molecular epidemiology in the genomic era (32). Our investigation fully supports the view that multisampling is critical to decipher transmission patterns, especially in the context of outbreaks with such long timeframes. Here, multiple sampling was only performed prospectively for four patients (P03, P04, P05 and P06) with one isolate stored and studied for all of the remaining patients. The proportion of mixed infection may thus have been higher than detected here if all of the isolates from all of the patients had been investigated. This work suggests that guidelines for sampling in future *Mucorales* outbreaks should be implemented.

Our management of this outbreak led to the implementation of a twice weekly screening of serum samples from patients hospitalized in BU1 using *Mucorales* PCR, and by immediate prescription of antifungal treatment when the PCR test was positive (25) in addition to isolation of all culture-and/or PCR-positive patient and dedicated nurses to prevent the risk of transmission to other patients. So far, with 2-years of hindsight, no additional case of *M. circinelloides* f. *circinelloides* IWM has been observed. This illustrates how hygiene prevention together with early diagnosis of mucormycosis may improve patient management and avoid dramatic outbreaks in hospital settings among populations at risk.

## MATERIALS AND METHODS

### Isolates and patients

An outbreak of *M. circinelloides* f. *circinelloides* infections was suspected in the Saint-Louis Hospital (SLS) located in Paris that involved six patients (P01 to P06) from BU1 between March 2013 and January 2015. The outbreak was not suspected until *M. circinelloides* was recovered from wounds of patient P04. Five of the six patients had proven IWM using a modified version of the EORTC/MSG criteria (25, 33) and one was considered colonized and was not treated.

In order to study the genetic relatedness between the BU1 clinical isolates, additional isolates identified as *M. circinelloides* from other sources were selected (Table 1): (i) sequential isolates from 4 of the patients prospectively collected from the skin lesions as previously described (25); (ii) isolates collected in SLS but in another ward (one from the environment, and another one colonizing a patient); (iii) isolates (n=4) recovered in another outbreak that involved 4 patients hospitalized in another burn unit (BU2) in the ‘Hopital d’instruction des Armées’, Clamart, located in the suburb of Paris (PER) over the same period of time. These isolates have been sent to the French National Reference Center for Invasive Mycoses & Antifungals (NRCMA); (iv) one isolate from invasive mucormycosis recovered from the kidney biopsy of a kidney transplant recipient in Strasbourg in the Eastern part of France (STR).

All positive slants were sub-cultured once on Sabouraud dextrose agar with gentamycin and chloramphenicol (Bio-Rad, Marnes-la-coquette, France) at 30°C using the bulk and never single colonies.

Overall, 21 isolates were selected for further analysis: 15 clinical isolates from BU1 (SLS) (12 bulk cultures including 1 from colonization), 1 environmental isolate from SLS, 4 clinical isolates from BU2 (PER), and 1 clinical isolate from STR.

The sequence of *Mucor circinelloides* 1006PhL strain was used as the reference genome (34).

### Environmental investigation in BU1

Extensive environmental sampling was performed and mycological contamination was investigated by culture methods on Sabouraud agar (Bio-Rad, Marnes-la-coquette, France) and 2% Malt extract incubated at 30°C and 37°C. Overall, 30 specimens from non-sterile material, air, surfaces, and aeration machineries, technical room, Bair Hugger machines, and dedicated non-sterile material were tested. Bair Hugger machine allows to actively warm the patient. It pulses warm air through a plastic pipe into a blanket applied on the patient.

### Polyphasic identification of isolates

The 21 isolates were sent to the NRCMA, where the purity was verified and identification to the species-level performed using phenotypic and molecular identification. In details, microscopic examination was performed on 5 to 7 day old cultures growth on 2% malt agar at 30°C. Amplification and sequencing of the ITS1-5.8S-ITS2 region and the D1/D2 region of the LSU rDNA were performed as described previously (35). The amplification of the RPB1 gene (RNA polymerase II largest subunit) was made with primers RPB1Ac and RPB1Cr (36). The PCR products were then sequenced and the consensus sequences were obtained as already described (35). Sequences were subjected to pairwise alignments against curated fungal reference databases available at the on-line MycoBank database (http://www.mycobank.org/).

### Sequence alignment and phylogenetic analysis

*Mucor circinelloides* is a single species consisting of four different formae (f. *circinelloides;* f. *griseocyanus;* f. *janssenii;* and f. *lusitanicus),* with formae *circinelloides* the most commonly involved in human mucormycosis (24). Multilocus sequence alignments on partial sequences of 3 different loci (ITS, 28S and RPB1) was performed (35, 37).

The sequences were aligned using MAFFT v.7.308 with default settings. Data from each gene was analyzed separately and combined as a concatenated 3-locus dataset. For the multilocus dataset, Neighbor-joining (NJ) phylogenetic trees were constructed using MEGA6 software (38) with Tamura 3-parameter substitution model and 1000 bootstrap replicates. The program PhyML v3.0.1 (39) was used to infer maximum likelihood (ML) phylogeny using TN93 substitution model and 1000 bootstrap repetitions. Bayesian analysis with default prior of Mrbayes v.3.2 (40) was conducted to determine posterior probabilities. Two analyses were done by running 10^6^ generations in four chains, sampling every 100 generations.

### Whole genome sequencing and assembly

Because genotyping methods for *M. circinelloides* f. *circinelloides* were lacking, WGS was performed to compare isolates. Libraries were constructed using a Nextera XT DNA sample preparation kit (Illumina) and sequenced with an Illumina NextSeq 500 sequencing system with a 2×150-nucleotide paired-end protocol. Two lanes from these tagged libraries resulted in ~12.6 million read pairs per strain on average.

All sequenced reads were clipped and trimmed with AlienTrimmer v.0.4.0 (41), corrected with Musket v.1.1 (42), merged (if needed) with FLASH v.1.2.11 (43), and subjected to a digital normalization procedure with khmer v.2.0 (44). For each sample, processed reads were finally assembled with SPAdes v.3.10 (45).

### Whole genome analysis

For each pair of assembled genomes, an evolutionary distance (proportion of aligned nucleotide differences) was estimated with Mash v.1.0.2 (sketch size = 100,000) (28) The resulting distance matrix was used to infer a Minimum Evolution phylogenetic tree with FastME v.2.1.5 (46, 47).

### Reproducibility of the sequencing process

In a specific experiment dedicated to determine the reproducibility of all the sequencing process (from extraction to sequence analysis), additional single spore isolation was performed on malt extract agar 2% plates for 3 isolates recovered from patients P03, P04 and P05 in BU1 (P05_600_BU1_SLS, P04_603_BU1_SLS, and P03_594_BU1_SLS, see Table 1). One colony was thus selected from each isolate for additional sequencing.

### Genome stability experiments

Genome stability during vegetative growths and host infections was analyzed with 1006PhL strain. In brief, the strain was grown on PDA for one day. The colony was streaked to isolate a single colony, which then transferred onto a new PDA. After four days of incubation at 30°C under the light, the spores were collected and subjected to the same procedures up to three passages (VEG1, VEG2 and VEG3 isolates).

For infection passages, spores were suspended in sterile PBS. Male 8 weeks-old BALB/c mice were infected with 10^6^ spores in 200 μL of sterile PBS via tail vein injection. At day 3 post-inoculation, the mice were sacrificed, their brain collected and placed onto PDA after brain homogeneization. After one day of incubation, the fungal colony emerging from the brain was streaked to isolate single colonies. One colony was transferred onto PDA and incubated at 30°C under light for four days. Spores were then used as inoculum for the next infection passage, up to three passages (INF1, INF2, and INF3 isolates). Genomic DNAs of these isolates were used to construct libraries, 180-bases fragments and 2 to 3 kb jumps, which were sequenced by an Illumina HiSeq 2000 platform at the University of North Carolina High-throughput Sequencing Facility (HTSF).

## Ethics Statement

The murine infection experiment was conducted at the Duke University Medical Center in full compliance with all of the guidelines of the Duke University Medical Center Institutional Animal Care and Use Committee (IACUC) and in full compliance with the United States Animal Welfare Act (Public Law 98-198). The Duke University Medical Center IACUC approved all of the vertebrate animal studies under protocol number A061-12-03. The studies were conducted in the Division of Laboratory Animal Resources (DLAR) facilities that are accredited by the Association for Assessment and Accreditation of Laboratory Animal Care (AAALAC).

## FUNDING INFORMATION

This study has been supported by Institut Pasteur and Santé Publique France. JH is supported by NIH/NIAID R37 MERIT Award AI39115-20, NIH/NIAID R01 Award AI050113-13, and Astellas Award CRES-15G01.

## ACKNOWLEDGMENTS

We thank Vincent Enouf from the P2M (Plateforme de microbiology mutualisée), Institut Pasteur, Paris for supervising and organizing the sequencing of the *M. circinelloides* f. *circinelloides* isolates.

